# Effects of selection stringency on the outcomes of directed evolution

**DOI:** 10.1101/2024.06.09.598029

**Authors:** Berk A. Alpay, Michael M. Desai

## Abstract

Directed evolution makes mutant lineages compete in climbing complicated sequence-function landscapes. Given this underlying complexity it is unclear how selection stringency, a ubiquitous parameter of directed evolution, impacts the outcome. Here we approach this question in terms of the fitnesses of the candidate variants at each round and the heterogeneity of their distributions of fitness effects. We show that even if the fittest mutant is most likely to yield the fittest mutants in the next round of selection, diversification can improve outcomes by sampling a larger variety of fitness effects. We find that heterogeneity in fitness effects between variants, larger population sizes, and evolution over a greater number of rounds all encourage diversification.

## Introduction

A common bioengineering goal is to create a protein that performs a specific function. One approach to this challenge is to use an existing protein as a template and apply biochemical reasoning to modify it such that it performs the new function [1]. An alternative and now widely used approach is directed evolution [2, 3], in which an experimenter starts from a template or set of templates, mutagenizes them randomly, selects from these mutants a new set of variants with improved function, and then repeats the process. Over multiple rounds, this process leads to the accumulation of multiple mutations that improve function, without the experimenter needing to characterize how they do so. In this sense, directed evolution is a step-by-step analog of natural selection, and has proved to be a powerful tool for bioengineering [2, 4] and understanding natural evolution [5]. Directed evolution has, for example, yielded enzymes more efficient than synthetic catalysts [6], experimentally useful fluorescence proteins [7], and insights into affinity maturation of broadly neutralizing antibodies [8]. In addition to proteins, this approach has also been used to engineer RNAs [9], synthetic genetic polymers [10], and genetic circuits [11].

A common way to implement directed evolution is to encode an initial protein sequence on a plasmid and then use error-prone PCR to create a plasmid library containing many variants of this initial protein [12]. This diverse library of protein variants is then transformed into a cellular display system [13], which couples the desired protein activity to cellular fluorescence in some way [14]. One can then use fluorescence activated cell sorting to select the best-performing cells [8, 15], extract the plasmids from these cells, perform another round of error-prone PCR, and repeat the process.

While this basic workflow — repeated mutagenesis and selection — is the core of any directed evolution approach, there are numerous possible variations [3]. For example, lower-throughput methods in which each protein variant is spatially separated [16] (e.g. across microplate wells) can provide more control of how they are selected and mutagenized. Other approaches trade control for speed and automation, for example in systems that allow cells to be selected via competition and mutagenized continuously [17, 18].

Regardless of the details of how the experimental workflow is implemented, directed evolution is blind to the underlying map from sequence to function [19]. Rather than using biochemical reasoning to choose the next sequences to test, the experimenter chooses the parameters of a population’s evolution. As with evolutionary adaptation in any system, the dynamics and outcomes of directed evolution depend on these choices. One key parameter is the mutation rate [20]. One wishes to generate variation that includes beneficial mutations, but not so much variation that these beneficial mutations are too often linked with and weighed down by deleterious mutations, which are typically more likely to occur [5]. Another choice is the number of mutants to generate at each round, which is typically determined by a tradeoff between practical constraints on the number of mutants that can be screened and the desire to have larger population sizes that help to discover more beneficial mutations at each round.

Here, we focus on a third key parameter [21] of directed evolution experiments: how stringently to select for improved function in each round. Selection stringency defines the likelihood that each variant is selected for mutagenesis in the next round (e.g. what defines the cutoff for “best-performing cells” in the workflow described above). For example, we might select the top half, top one percent, or even just the single best variant at each round. Or alternatively, we might select in some more complex way, for example by seeding the next round primarily with mutants of the fittest variant in the previous round, but also with a smaller number of mutants of less-fit variants. On the one hand, we must impose some form of selection or there would be no pressure for variants to climb to greater fitness through successive rounds. On the other hand, imposing too harsh a selection pressure limits our ability to explore the sequence-function landscape, and could potentially lead to the process becoming trapped at a local optimum. The optimal choice of selection stringency is unclear, but it must involve some balance between greedy exploitation of the fittest variants versus a more relaxed selection that allows for broader exploration of the landscape [22].

Work in adjacent fields has developed a variety of approaches to this question. For example, in computer science, active learning approaches integrate available sequence-function data to create a computational model of the landscape that is then used to choose the set of sequences to screen at the next round in a way that will optimize fitness gains while gaining additional information about the landscape [23, 24]. These methods can be powerful and efficient, but they rely on high-throughput direct measurements of sequence-function relationships, along with the construction of custom libraries of specific chosen variants. Instead, we consider here the simpler approach of directed evolution by random mutagenesis. This is analogous to analysis of the short-term effects of selection stringency in population genetics, which have historically been studied in the context of plant and animal breeding [25], considering mediating factors such as heritability, inbreeding, and frequencies of standing variants [26]. More recently, studies of protein evolution have described responses to selection stringency [20, 22, 27] and the biophysical mechanisms explaining different outcomes [28] in specific systems.

Here, we focus instead on the general problem of how the structure of the sequence-function landscape affects the optimal choice of selection stringency. In practical experimental settings, there is noise in the measurement of function and therefore in our estimates of the relative fitness of each variant. However, we will consider the idealized case in which measurement noise can be neglected, and instead characterize the optimal selection when the exact fitness of each variant is known. One straightforward strategy, especially in this idealized case, is to select only the single fittest variant at each round. This strategy is optimal if the sequence-function landscape is perfectly smooth, meaning there are no non-additive interactions between the fitness effects of different mutations (i.e. no epistasis). On a perfectly smooth landscape, a given mutation will have the same effect on function regardless of which variant it occurs in, so greedily selecting the fittest variant at each round will tend to yield the fastest improvement in function. On the other hand, if the sequence-function landscape is rugged, the effect of a given mutation can vary greatly by sequence context. The magnitude of improvements available to different variants can vary dramatically and the fittest variant is not guaranteed to have the best evolutionary prospects. In this case, greedily selecting the fittest variant in each round may not be optimal, and less stringent selection that allows for more exploration of the landscape may be preferable.

This reasoning suggests that the ruggedness of the landscape is critical to determining the optimal selection stringency. Extensive prior work has attempted to quantify this ruggedness by empirically characterizing protein sequence-function landscapes. Broadly speaking, much of this work finds that epistasis is widespread and that sequence-function landscapes are at least to some degree rugged [29, 30]. For example, studies have created combinatorially complete libraries that consist of all possible combinations of some set of mutations separating two variants of a protein [31–34]. This work has shown that there are numerous “idiosyncratic” epistatic interactions between specific mutations, which constrain the potential trajectories that evolution could have taken. Other studies have assayed the effects of libraries of specific mutations on different ancestral sequences, again typically finding numerous epistatic interactions between the background sequence and mutational effects [35, 36]. However, there are also counter-examples [33], and the complexity of protein sequence-function landscapes remains controversial [37, 38]. Thus the overall extent to which epistasis creates ruggedness in protein sequence-function landscapes, and how this ruggedness affects the optimal selection stringency in directed evolution, remains unclear.

An alternative body of work has used theoretical models of fitness landscapes to explore how selection stringency and other parameters affect the statistics of evolutionary trajectories. For example, extensive work has analyzed adaptive walks in the NK model [39, 40], which parameterizes the landscape in terms of the number of epistastic interactions each locus participates in. Other work has analyzed evolutionary dynamics in numerous other types of theoretical landscapes [41]. These landscape models are typically parameterized in terms of some set of genetic loci, their effects, and the epistatic interactions between them. In other words, they generate the landscape “microscopically” [42], in terms of specific epistatic interactions between particular loci. An alternative class of models are defined geometrically (e.g. Fisher’s geometric model [43] or the Rough Mount Fuji [44] models), or relatedly based on phenotypic correlations that decay with genetic distance (e.g. [45]).

In principle, one can use these existing theoretical landscape models as the basis for investigating the effects of selection stringency on the dynamics and outcomes of a directed evolution experiment (see e.g. recent work focusing on the NK model [22]). However, the effect of selection stringency does not depend on all of the complex details of microscopic epistasis or the full geometric structure of the landscape. Instead, the key question is how the spectrum of potential adaptive mutations varies across different genetic backgrounds. In other words, how does the accumulation of one mutation (or a combination of mutations) change the *distribution of fitness effects* (DFE) of potential future adaptive mutations? This effect has been termed “macroscopic” epistasis [42]. While macroscopic epistasis ultimately arises from the collective effects of many microscopic interactions, the effects of ruggedness on the dynamics of directed evolution are more clearly described in terms of the former.

Motivated by this, we investigate here the effects of selection stringency on directed evolution in several simple models of macroscopic epistasis. Specifically, we imagine that each variant has some DFE which is in some way changed by mutation. There are many possible models of these changes in the DFE, including for example the general pattern of diminishing returns epistasis that has been observed experimentally in several systems [46–48]. However, while this form of macroscopic epistasis leads to a systematic trend of declining adaptability as fitness increases, it does not lead to ruggedness that strongly favors exploration in directed evolution, because all equally fit sequences suffer equally. We therefore focus instead on other, more rugged patterns of macroscopic epistasis, in which mutations idiosyncratically change DFEs. We measure how the optimal selection stringency for directed evolution depends on the ruggedness of the model of macroscopic epistasis, as quantified by the heterogeneity in the DFEs of candidate variants. We begin in the next section by analyzing a toy model of selection among two variants in a single round of a directed evolution experiment. We then expand this model in subsequent sections to analyze selection among an arbitrary number of variants and over multiple rounds of directed evolution.

## Results

### Diversification can help explore heterogeneous DFEs

We begin by imagining that we have a set of variants (either our starting library or the variants generated from a previous round of directed evolution) and we now need to select the ones that pass the selection threshold and serve as the basis for mutagenesis in the next round of directed evolution. Among this set of variants, one of them is the fittest. Since it already has the most successful sequence, it is natural to ask: why not simply select only this one? In other words, why not impose the maximum possible stringency of selection? In this section, we will ask why it might be favorable to adopt a less stringent selection pressure, and instead select a more diverse pool of variants. We do so in the context of a toy scenario, in which we select among only two variants of different fitness, to illustrate the essential tension between exploration and exploitation of the sequence-function landscape (Fig. 1A).

**Figure 1.**
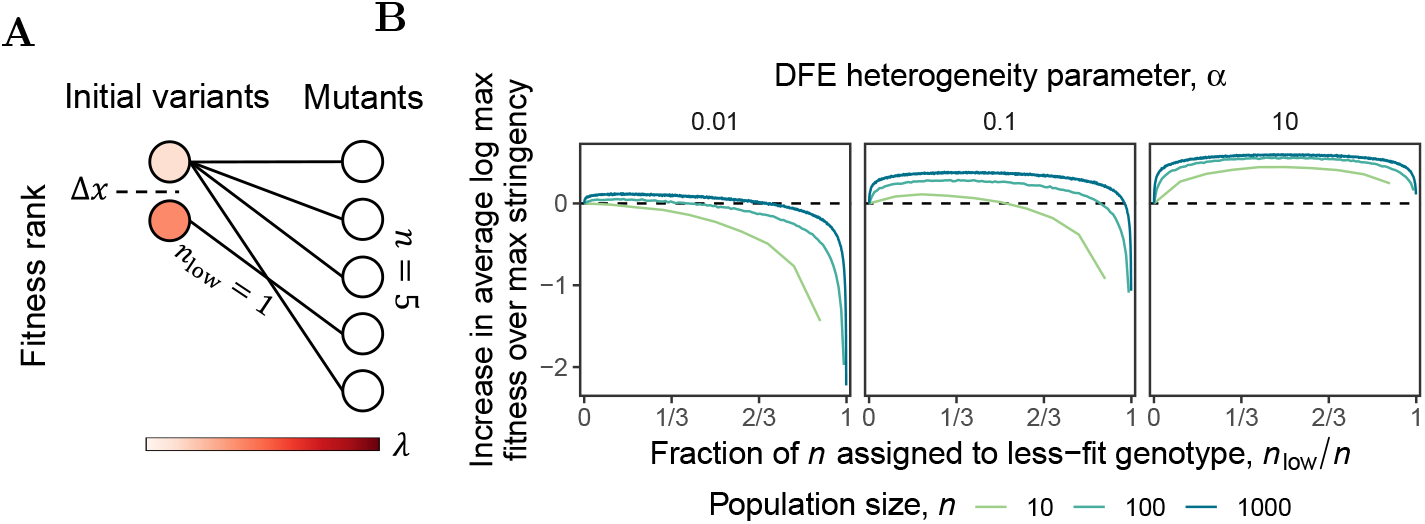
The immediate effect of selection stringency on fitness while drawing mutants from two parents. **(A)** Schematic of our two-parent model, in which we assume that we generate mutants for the next round (here a total of *n* = 5 mutants) from two parental variants of different fitness. In this model, the selection stringency is determined by the number of mutants we draw from the lower-fitness parent, which has a fitness disadvantage of Δ*x* compared to the higher-fitness parent. We assume that each parent has an exponential DFE, with parameter *λ* that is drawn at random as described in the text. **(B)** Simulations of the two-parent model (with Δ*x* = 1) showing how the maximum fitness of the mutants depends on the selection stringency, i.e. the fraction of the mutants drawn from the less-fit parent. Note that the advantage of diversification increases with *n* and with the degree of heterogeneity between the DFEs of the parents.

Specifically, starting from these two variants, imagine we are limited by experimental constraints to construct and screen a total of *n* mutants in the next round. However, we can decide how many mutants of each variant will compose that screen. The question is thus how many mutants to “draw” from each variant. Imagine that we take *n*_high_ mutants of the fittest variant and the remaining *n*_low_ = *n* − *n*_high_ mutants from the less-fit variant. Our goal is to understand how the fitness of the fittest variant in the next round depends on *n*_low_. If this next-round fitness is maximized with *n*_low_ = 0, then maximal selection stringency is preferred, i.e. only the most-fit variant is selected for mutagenesis in the next round. However, if some *n*_low_ *>* 0 is better, then it means that less stringency is preferred (up to a maximum of *n*_low_ = *n/*2, which means that the two variants are equally mutagenized and hence corresponds to the largest possible diversification in this toy model).

It is natural to suppose that mutants of higher-fitness variants tend on average to be more fit than mutants of lower-fitness variants. Thus at first glance it may appear that choosing *n*_low_ = 0 (i.e. maximal selection stringency in which we mutagenize only the fittest variant) might be optimal. However, it is possible that in some cases mutations on the background of the most-fit variant are less favorable than on that of the less-fit variant. For example, two proteins of similar fitness may differ greatly in evolvability if one is quite stable and the other only marginally so [49]. This difference in stability could arise for example due to some apparently neutral mutation [50]). Indeed, it is often the apparently neutral (from the standpoint of the fitness assay) but stabilizing mutations that go on to enable performance of a function [50]. While stability is likely to be a key molecular phenotype underlying evolvability, other molecular phenotypes [51] such as the structural organization of the protein fold [52] and conformational diversity [53] might also play similar roles.

Regardless of the origins of differences in evolvability, the important point is that even a small number of mutations can significantly alter the effects of other mutations and therefore the evolutionary prospects of variants [54, 55]. Thus, even if on average mutants of higher-fitness variants tend to be more fit, the opposite can also sometimes be true. Even if this is only rarely the case, it can be advantageous to devote some resources to mutagenizing the lower-fitness variant as well. The extent to which this is true (and hence the best choice of *n*_low_) will depend on how often and to what degree genetic backgrounds differ in their favorability to mutation.

To analyze this situation of selecting among two starting variants more quantitatively, we introduce a simple toy *two-parent model*) (Fig 1A, Methods). By definition, the fittest variant has greater fitness than the less-fit variant. We will assume this fitness difference is Δ*x*. The larger Δ*x*, the greater the advantage of sampling the fittest variant; that is, the greater the advantage of exploitation. However, we assume that the distributions of fitness effects (DFEs) available to the two variants can also differ. For the sake of concreteness, we assume that the DFE of beneficial mutations for both variants are exponential, with rate (i.e. inverse scale) parameters *λ*_low_ and *λ*_high_, respectively. Thus the fitnesses 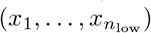 of the mutants of the lower-fitness variants are drawn independently as Exponential(*λ*_low_) − Δ*x*, while the mutants of the higher-fitness variants 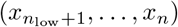 are drawn independently as Exponential(*λ*_high_). The outcome we measure is the resulting maximum fitness of the new population of *n* mutants, *M* = max(*x*_1_, …, *x*_*n*_).

If *λ*_low_ = *λ*_high_, there would be no advantage to diversification, as the two variants have the same DFE but the higher-fitness variant enjoys an initial fitness advantage. In this case *n*_low_ = 0 clearly maximizes *M*. However, suppose that there is some random variation in *λ* between the variants. Even if *on average λ*_low_ = *λ*_high_, we will sometimes have *λ*_low_ *< λ*_high_. This can create an advantage to diversification, such that *n*_low_ *>* 0 maximizes *M*. The key point is that there are diminishing returns to drawing more mutants from a single exponential distribution. This can make it advantageous to draw fewer mutants from the most-fit variant and instead devote some resources to sampling from a second DFE with a different *λ*, even if this comes at the price of starting at an initially lower fitness. Whether this is true will depend on the typical scale of variation in *λ* (which determines the potential advantage of sampling from a second DFE), the difference in fitness between the variants Δ*x* (which determines the penalty we pay for starting from a less-fit variant), and the total number of the mutants we are screening *n* (which determines the extent to which there are diminishing returns to drawing additional mutations from the first DFE).

To illustrate this point, we can quantify the variation in DFEs by assuming that *λ*_low_ and *λ*_high_ are themselves random. Specifically, we consider the case where they are independently drawn from exponential distributions with parameter *α*. Here *α* parameterizes the degree to which DFEs tend to differ between variants: the mean effect of a beneficial mutation (and the standard deviation of these effects) has an interquartile range of approximately 2.8*α* (SI Appendix). In other words, the heterogeneity of the DFEs increases with *α* (and because the *λ* are drawn independently, each variant is equally likely to have the more favorable DFE).

In Fig. 1B, we show how the maximum fitness of the mutants in the next round depends on the degree of diversification *n*_low_ for several different values of *α* and *n* (note that as an extreme value statistic, the convergence of the expected maximum *M* is sensitive to model details, so we instead plot how the expected log *M* depends on these parameters). For sampling mutants from the lower-fitness variant to be advantageous, its DFE must be more favorable to such a degree that it overcomes its Δ*x* fitness disadvantage. This becomes more likely as *α* becomes larger relative to Δ*x*. Thus as *α* increases, the optimal number of samples to draw from the less-fit variant also increases up to the point of maximal diversification, *n*_low_ = *n/*2 (because it is equally likely that the DFE of the more-fit parent is more or less favorable than the DFE of the less-fit parent, it is never optimal to increase *n*_low_ beyond this point).

The optimal number of samples to draw from the less-fit variant also depends on the number of mutants that can be screened, *n*. Particularly for larger *n*, it is optimal to diversify somewhat (i.e. the optimal *n*_low_ *>* 0) even when *α* is small compared to Δ*x*. In these cases, it is very unlikely that the DFE of the less-fit variant is sufficiently more favorable than the DFE of the more-fit variant to overcome its initial fitness disadvantage. Nevertheless, because of the diminishing returns of continuing to sample more mutants from the DFE of the more-fit variant, given sufficient *n* and *α* it is still optimal to spare some samples for a second DFE: the chance this DFE is anomolously favorable is larger than the chance that an additional sample from the DFE of the more-fit variant will be more fit than all previous samples. For this reason, for sufficiently large *n* it can even be optimal to favor maximal diversification (i.e. *n*_low_ = *n/*2) even when *α* is small compared to Δ*x*.

We can quantify this effect by calculating the *α*^*^ at which it becomes advantageous to reduce stringency to *n*_low_ = *n/*2. We can think of this threshold *α*^*^ as an upper bound on the *α* that would justify more moderate diversification. When *n* = 2, a straightforward calculation of the expected *M* as a function of *n*_low_ shows that *α*^*^ = Δ*x* (SI Appendix). Thus, when sampling a small number of mutants from two parents, the variation in their DFEs must be on the order of the fitness differences between them to justify diversification. However, as we can see in Fig. 2, *α*^*^ rapidly declines as *n* increases (e.g. *α*^*^ = Δ*x* when *n* = 2 but decreases to *α*^*^ ≈ 0.29Δ*x* at just *n* = 4).

**Figure 2.**
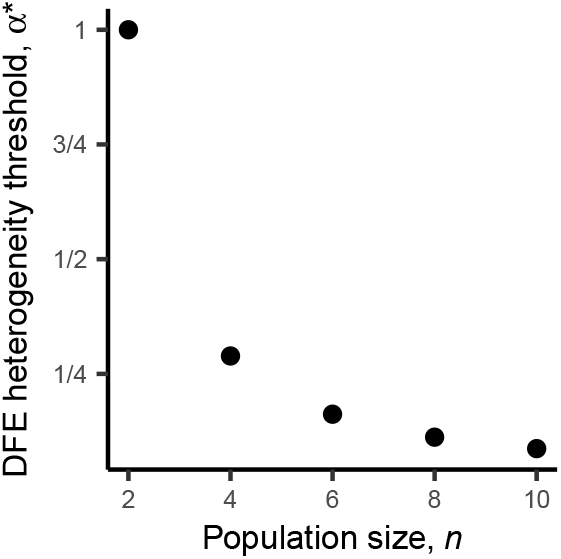
DFE heterogeneity required to justify maximal diversification. The *α*^*^ at which it becomes more advantageous to diversify maximally than to not diversify at all in our two-parent model (where the DFE rate parameters *λ* Are distributed as Exponential(*α*)). Note that, for small *n*, the DFE heterogeneity *α* must be roughly on the order of the fitness difference between parents Δ*x* to justify maximum diversification. However, as *n* increases, maximal diversification can be favorable even when *α* is substantially less than Δ*x*.

### The benefits of sampling a wider range of DFEs

Thus far we have analyzed the effects of diversification in the context of a simple toy model involving only two parental variants. In this model, we could quantify the degree of selection stringency entirely in terms of the fraction of *n* mutants that are drawn from the less-fit parent. However, in practice we typically have more flexibility: if we screen a total of *n* mutants in a given round of directed evolution, we can select any subset of these as parents for the next round.

In this section, we consider this more general case. Specifically, we imagine that out of the *n* mutants in the current round, we select the most-fit *k* variants as parents for the next round (since we will assume the DFE of each variant is drawn independently, there is never an advantage to omitting some of the fitter variants in favor of less-fit ones). In principle we could imagine that mutants for the next round are drawn in some complex way from these *k* parental variants. However, for simplicity and concreteness (and consistent with the practical constraints of many directed evolution workflows), we imagine that we draw mutants for the next round about equally from each of these parents for a total of *n* variants in the next round. Our goal is to understand how the maximum fitness, *M*, of these *n* total variants depends on *k*. If *k* = 1 is optimal, we should maximize selection stringency by drawing all mutants from the most-fit variant in the current round. If on the other hand *M* is maximized for some *k >* 1, then at least some diversification is favorable, up to the maximal possible diversification of *k* = *n*.

We illustrate this scenario, which we call the *k*-parent model, in Fig. 3A. As in the two-parent model, we assume that the DFE of each parent is exponential with some parameter *λ*. As before, we model random heterogeneity in the DFEs by assuming that *λ* is drawn independently for each parent from an exponential distribution with parameter *α*, so larger *α* corresponds to greater heterogeneity in DFEs between variants.

**Figure 3.**
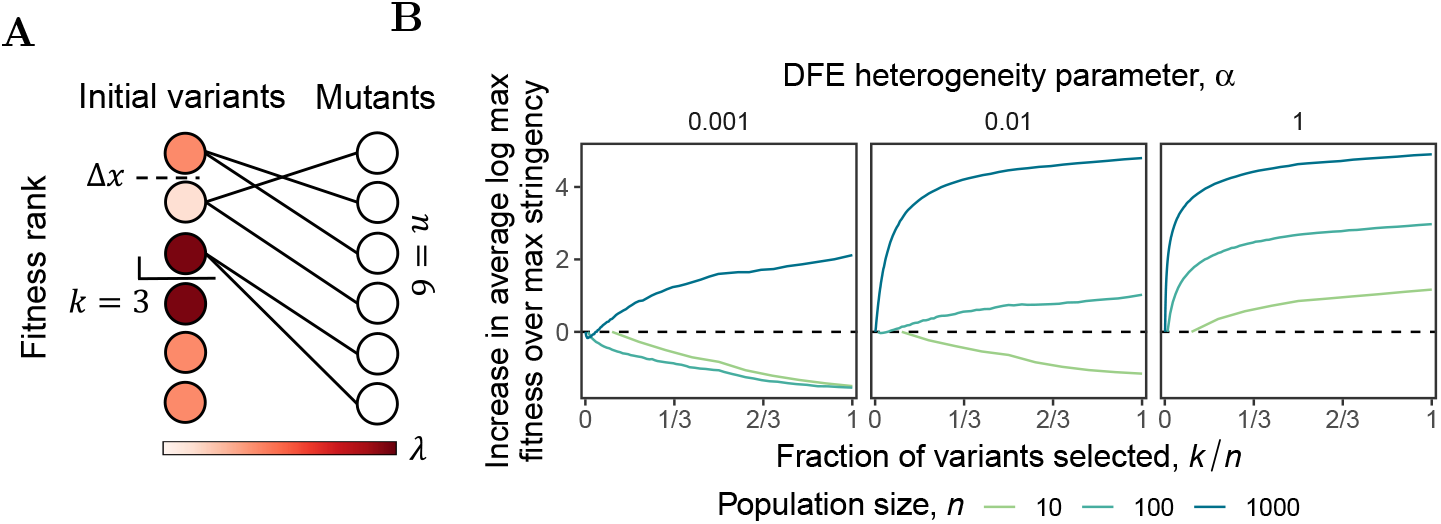
The effect of selection stringency in the *k*-parent model. **(A)** Schematic of the *k*-parent model, here with *n* = 6 and *k* = 3. **(B)** Simulations of the *k*-parent model (with Δ*x* = 1) showing how the maximum fitness of the mutants depends on the selection stringency, i.e. the fraction of variants selected as parents for the next round, *k/n*. Note that the advantage of diversification increases with *n* and with the degree of heterogeneity between the DFEs of the parents.

We could make a variety of assumptions about the relative fitness differences between the *k* parental variants. For simplicity, we assume here that the most-fit variant is Δ*x* fitter than the other *k* − 1 variants, which are of equal fitness. This choice ensures that the benefit of diversifying among *k* variants that we observe will be a lower bound on the true benefit in a more complex model in which Δ*x* is the difference between the fittest and least-fit of the *k* variants (with the other variants intermediate between them). Of course, in practice the value of Δ*x* will tend to increase with *k*, so we can also interpret Δ*x* as determining the best choice of *k*, by setting the bound on the extent to which we want to diversify among less-fit variants.

In Fig. 3B, we show how the maximum fitness of the mutants in the next round depends on the choice of selection stringency *k*, for several different values of *n* and *α*. Our results in this *k*-parent model are qualitatively similar to those from the two-parent model: the advantage of diversification increases with the number of mutants, *n*, and with the degree of heterogeneity between the parental DFEs, *α*. They are qualitatively similar for the same essential reason: there is diminishing returns from sampling many mutants from a single DFE, so provided that the total number of mutants and the degree of heterogeneity between DFEs are sufficiently large, the cost of starting from less-fit parents is outweighed by the advantage of sampling from more than one DFE. We also note that this result is not specific to the details of how we model DFEs and the heterogeneity between them; the same basic dynamic is recapitulated in a model where DFEs are normally distributed and *α* controls the distribution of their means (Fig S1).

### Inherited changes in DFEs drive the value of diversification

Thus far, we have analyzed the effects of selection stringency on the maximum fitness of a set of mutants in a single round of directed evolution. If we assume that the dynamics at each round are identical and independent, then the optimal selection stringency across multiple rounds of directed evolution should simply be repeated use of the optimal single-round stringency. However, it may often be the case that variation in DFEs is not independent across multiple rounds. For example, if a particular protein variant has a less-favorable DFE because it is barely stable, most of its descendants are also likely to be barely stable and hence also have less-favorable DFEs, and vice versa [49]. In other words, among proteins of similar fitness, the DFEs of more stable proteins can be expected not only to be superior in the current round of evolution, but one might also expect the DFEs of their mutants to be superior than the DFEs of mutants of less stable proteins.

To analyze these effects of heritability in DFEs, which violate the assumption that the dynamics at each round are identical and independent, we consider here an extension of our *k*-parent model. In this *k*-parent *inheritance* model, each variant continues to have two properties: a current fitness and a DFE for mutations in the subsequent round. However, we now assume that the DFE parameters *λ* for each variant are not drawn at random in each round, and instead are inherited (but imperfectly, to maintain some model of the generation of heterogeneity). Specifically, rather than being drawn at random for each variant, we assume that all variants initially have identical DFEs, which are inherited by their offspring. However, each DFE has some constant probability of becoming heritably less favorable at each round (for example because that particular adaptive mutation has a destabilizing effect on the protein). In this setting, maintaining some diversity at each round can be beneficial not only in maximizing fitness in the immediately following round, but also in maintaining high-evolvability lineages that promote adaptation in the future.

To implement this model, we assume that all variants have an exponential DFE with scale parameter *β*, which is initially identical for all variants. At each round, mutants inherit their parental DFE, with some chance *p* that their *β* decreases by an amount *d* (though because the exponential distribution must have positive scale, we set a minimum *β* of 1*/*100; once this limit is reached the DFE cannot continue to degrade). Subsequent selection is performed similarly to the *k*-parent model, and we quantify the selection stringency using the parameter *k* as before. We find that the value of diversification generally increases as the probability of degradation of the DFE goes up or as we consider directed evolution across a larger number of rounds (Fig. 4). Because these two factors control the extent of heterogeneity in the DFEs, this is analogous to our results from the two-parent and *k*-parent models (though we note that the dependence on the number of rounds suggests that, instead of enforcing a constant selection stringency *k*, decreasing *k* at each round would improve the outcome).

**Figure 4.**
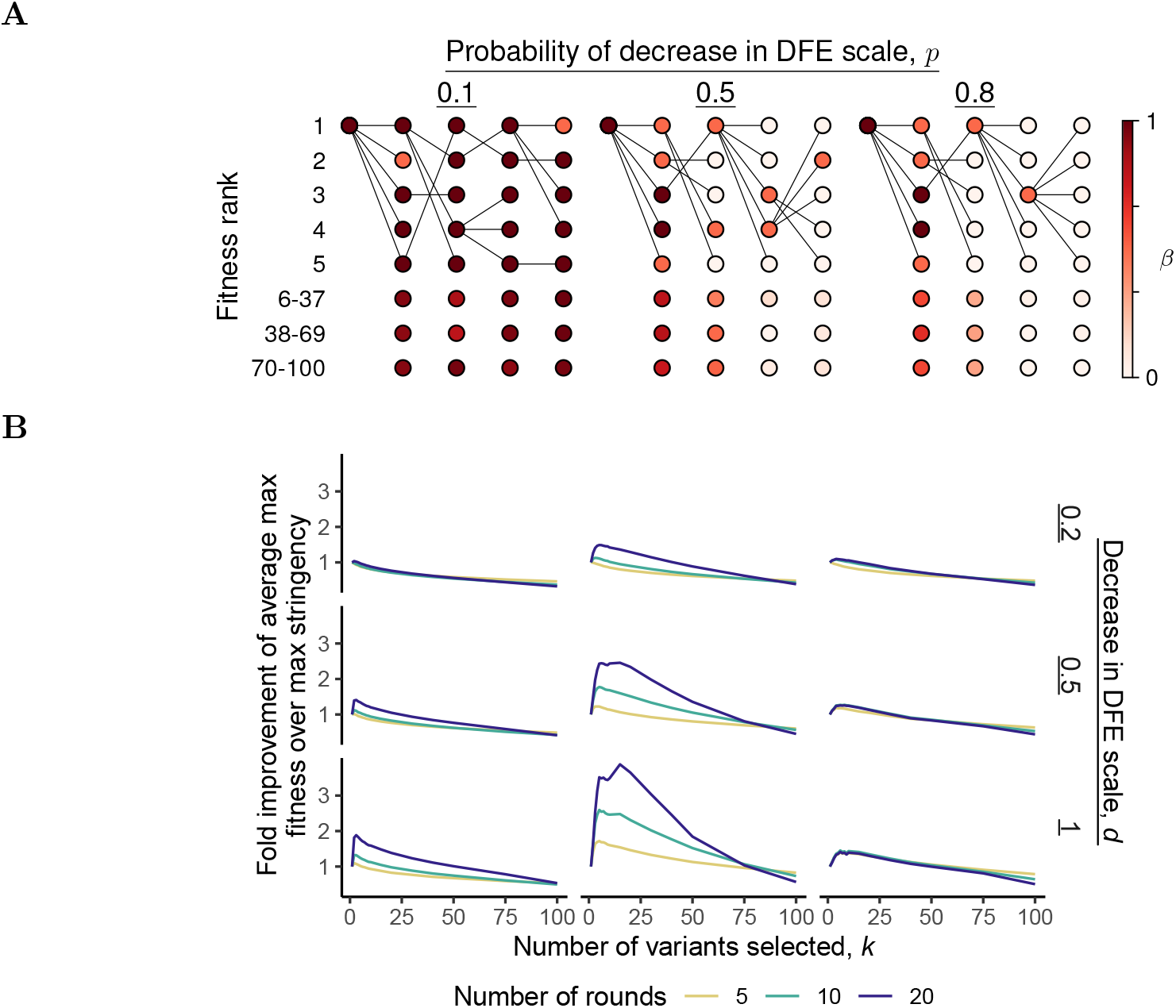
Effects of selection stringency in the *k*-parent inheritance model. **(A)** Examples of the dynamics across four rounds of directed evolution in our *k*-parent inheritance model for three different probabilities *p* of DFE degradation. Here *d* = 0.5, *n* = 100, and the selection stringency is *k* = 5. For variants not among the fittest *k*, color reflects the average *β* in that bin. Edges indicate parentage, though edges to variants below the top *k* are not shown. **(B)** At a population size of *n* = 100, the average maximum fitness at the end of 5, 10, and 20 rounds of directed evolution as a function of selection stringency *k* and the features of DFE inheritance (*p* and *d*).

In Fig. 4A, we show specific examples of evolution at different rates of DFE degradation to illustrate these general trends. When *p* is low, occasional lineages with unfavorable DFEs are effectively purged, and several of the top variants in each round are viable because of the stability of DFEs. However, at moderate *p*, only some lineages remain viable and the fittest variant at the end of the evolution has an ancestry widely ranging in fitness rank, not being descended from the fittest variant at each round. This indicates that maintaining evolvability of the DFE plays a critical role, making it important to retain several candidate variants at each round. Finally, at even higher *p*, diversification is able to retain the most evolvable lineages for a short time, but DFEs quickly deteriorate to a minimum and diversification becomes futile. This reflects the fact that in our model DFEs become monotonically less favorable over time. If we allow for some small probability that mutations can occasionally improve the DFE (e.g. by improving stability [49, 56]), diversification can also be favorable because it helps to cast a larger net for mutations that improve evolvability (Fig S2). If *k* = 1, for example, there is only one chance in each round for the parental DFE to have improved. In contrast, at the cost of retaining some variants of comparatively lower fitness and unfavorable DFEs, diversifying increases the probability of a DFE improvement that can underlie fitness improvements over several subsequent rounds.

## Discussion

In directed evolution, some number of mutants can be screened at each round. Mutants modify the genetic sequence of their parental variants, thus exploring their DFEs — the effects of mutations on those backgrounds. Since there is a dropoff in fitness between the fittest variant and the rest, if the variants were equally evolvable, there should be no advantage to selecting any variant other than the fittest. By selecting less-fit variants, one would only sacrifice samples that could be used to better exploit the genetic background of the fittest variant. However, if the statistics of adaptation in the landscape around variants vary, reflecting idiosyncratic patterns of macroscopic epistasis, then there can be an advantage to diversifying. The advantage depends on the form of this variance and the number of mutants available to sample it. Our results show that as the heterogeneity in the DFEs relative to the magnitude of the fitness dropoff increases, so does the value of diversification. At sufficiently large population sizes, the DFEs of less-fit variants can be explored with the expectation that some of their mutants will often be fitter than those of the fittest ones. Imperfect heritability of DFEs leads naturally to this heterogeneity, with greater risks to the favorability of a DFE calling for more diversification.

In more realistic settings, there may be additional, higher-order heterogeneities in evolvability than are reflected in our simple models. For example, in our multi-round model (the *k*-parent inheritance model) we assume that DFEs are inherited variably, either perfectly or imperfectly. The probability of imperfect inheritance and the corresponding magnitude of the DFE degradation were assumed the same across all variants. However, one can imagine that a mutation on a particular background could, for example, impact not only the DFE but the rate and magnitude of future mutational effects on DFEs. Indeed, theoretical modeling of fitness landscapes has shown that the evolutionary history of two sequences can be an important differentiating factor of the evolvability of two sequences, even if their DFEs are similar [45]. The presence of such higher-order heterogeneities would seem to encourage greater diversification.

Throughout our analysis, we assumed DFEs are exponentially distributed, consistent with many previous theoretical studies which model beneficial fitness effects [57, 58]. The empirical evidence for exponential fitness effects of beneficial mutations is mixed [59], as is the evidence for whether the theoretical conditions underlying the exponential assumption [60] are satisfied [5, 61, 62]. In assuming exponentially distributed fitness effects, we also assumed that all effects are beneficial. We chose to focus on beneficial effects since they are the ones that drive directed evolution; mutants are typically not selected if they do not improve. However, although we made these assumptions for concreteness and tractability, we believe our general conclusions are robust to the specific choice of DFE model. For instance, we found that we could reproduce a core set of results with an alternative model relying on a different set of distributional assumptions (Fig S1). However, quantitative interpretation of the parameters and results will vary from experiment to experiment. For example, typically fitness effects are mostly deleterious [5, 63]. One should therefore interpret the population size parameter *n* as a fraction of a larger population size that also contains deleterious mutants. In proteins, this fraction is likely to be small (e.g. less than 1% [5]).

Throughout, we have also implicitly assumed a fixed mutation rate. As the mutation rate increases, many beneficial mutations may become linked to deleterious ones, leading to an effective change in the DFE. The balance between beneficial and deleterious mutations in such a setting will depend on the structure of epistasis and the set of mutants that happen to be generated. For example, it has been observed that high mutation rate causes greater variance in outcomes, sometimes leading to superior outcomes while risking inferior ones [20]. Future theoretical work could consider the role of this critical parameter on the course of directed evolution more generally.

While our results help quantify how optimal selection stringency depends on patterns of idiosyncratic macroscopic epistasis, it is less clear what these patterns are in any specific setting. Some inferences can be drawn from previous observations of the effects of selection stringency. For example, in prior simulated [27] and experimental [20, 21, 64] work, high stringency has typically corresponded to better outcomes than low stringency, indicating that DFEs between competitive variants were not consistently of great heterogeneity. The advantage of some degree of diversification has, however, also been recognized. For example, it has been observed that extreme stringency “is likely to be detrimental” [27], and suggested that low stringency at a low mutation rate might be useful in early rounds to produce diverse, viable variants [20].

To apply this work to design and optimize directed evolution experiments, the heterogeneity of DFEs during adaptation must be better understood. Such information should be easier to derive experimentally than sequence-function maps, as only the distribution of the phenotype of interest need be measured, as opposed to paired data consisting of the phenotype of each sequence. The heterogeneity will depend on, among other factors, the particular protein and the assay, but at least such experiments would sketch the possible range of heterogeneity and may indicate general behavior of DFEs over protein evolution.

## Methods

Code can be found at https://github.com/berkalpay/direvo-stringency. Average maximum fitnesses for each parameter setting were computed using 100,000 samples for the two-parent and *k*-parent models, and 2,000 samples for the *k*-parent inheritance model. We describe the three main models used in this paper below.

### Two-parent model

In the two-parent model, we assume there are two variants with a Δ*x >* 0 fitness difference between them, the fitter parent of fitness 0 and the other of fitness −Δ*x*. We assume a total of *n* mutants can be screened in the next round of directed evolution, *n*_low_ mutants drawn from the less-fit parent and the remaining *n* − *n*_low_ drawn from the fitter parent. We consider only the effects of beneficial mutations, which are drawn from exponential DFEs: the DFE of the fitter parent is Exponential(*λ*_high_) while that of the less-fit parent is Exponential(*λ*_low_). We assume that the DFE rate parameters are themselves random and drawn as 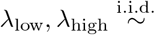 Exponential(*α*).

### *k*-parent model

Our *k*-parent model extends the two-parent model. We now assume there are *n* initial variants with a fitness difference of Δ*x* between the fittest and all of the *n* − 1 remaining variants. We consider a single round of directed evolution, in which we select the top *k* parents (the fittest and a random subset of the *k* − 1 remaining variants). Each parent gives rise to *ln/kj* mutants, with the remaining *n* mod *k* mutants assigned randomly without replacement. Mutant fitnesses are calculated as in the two-parent model, with each DFE being determined by a rate parameter drawn i.i.d. as Exponential(*α*).

### *k*-parent inheritance model

The *k*-parent inheritance model extends the *k*-parent model, with mutants becoming parents in the following round of directed evolution. We assume that the initial population is seeded by a single variant which has an exponential DFE with scale parameter *β* = 1. This initial variant is mutagenized to yield the starting population of *n* variants. At this and all subsequent rounds, a mutant inherits its parental scale parameter except with a certain probability *p* that its DFE scale *β* decreases by *d* (down to a minimum possible scale parameter of *β* = 1*/*100). At each round, we select the fittest *k* variants, and draw mutants from among these variants, with the number drawn from each parent multinomially distributed with equal sampling probabilities. In the illustrated examples, the number of mutants is divided equally between the parents.

## Acknowledgements

We thank Thomas Dupic, Jason Yu, Caelan Brooks, Caroline Holmes, Jeffrey Chang, Andrew Murray, and Angela Phillips for discussions and helpful comments. BAA acknowledges support from the NSF Graduate Research Fellowship Program (DGE-2140743). MMD acknowledges support from grant PHY-1914916 from the NSF and grant GM104239 from the NIH.

