## Supplemental Information for "Effects of selection stringency on the outcomes of directed evolution"

### SI Appendix for "Effects of selection stringency on the outcomes of directed evolution"

#### DFE properties under the two-parent model

The mean and standard deviation of an exponential DFE is the inverse of its rate parameter $\lambda$. In the two-parent model, this quantity $\beta = 1/\lambda$ is a random variable where $\lambda \sim \text{Exponential}(\alpha)$. Performing a transformation of variables, we have that

$$f_\beta(\beta) = \left|\frac{d\lambda}{d\beta}\right| f_\lambda(1/\beta) = \frac{\alpha}{\beta^2} e^{-\alpha/\beta}.$$

It follows that the distribution function is $F_\beta(x) = e^{-\alpha/x}$ and the quantile function is $Q_\beta(p) = \alpha/\log(1/p)$. The interquartile range is then

$$Q_\beta(3/4) - Q_\beta(1/4) = \frac{\alpha \log 3}{\log \frac{16}{9} \log 2} \approx 2.8\alpha.$$

The mode of the distribution is $\alpha/2$, the $\beta$ at which $f'_\beta(\beta) = \alpha e^{-\alpha/\beta}(\alpha - 2\beta)/\beta^4 = 0$.

#### Mutant fitness under the two-parent model

In the main text, we analyze statistics of the expected log maximum mutant fitness. For ease of notation, let us abbreviate $\lambda_{\text{low}}$ as $\lambda_{\text{L}}$, $\lambda_{\text{high}}$ as $\lambda_{\text{H}}$, and $n_{\text{low}}$ as $n_{\text{L}}$. The density of $M = \max\{x_1, \ldots, x_n\}$, the fitness of the fittest mutant, can be expressed as

$$p(M = m) = \int_0^\infty \int_0^\infty p(M = m|\lambda_{\text{L}}, \lambda_{\text{H}}) p(\lambda_{\text{L}}) p(\lambda_{\text{H}}) d\lambda_{\text{L}} d\lambda_{\text{H}}.$$

There are two cases: the fittest mutant is a mutant of the fittest variant, i.e. $m \in \{x_1, \ldots, x_{n_{\text{L}}}\}$, or it is a mutant of the other variant, i.e. $m \in \{x_{n_{\text{L}}+1}, \ldots, x_n\}$. Call the first event $L$ and the second $H$. Then

$$p(M = m|\lambda_{\text{L}}, \lambda_{\text{H}}) = p(M = m|\lambda_{\text{L}}, L)p(L) + p(M = m|\lambda_{\text{H}}, H)p(H).$$

$L$ means that all the mutants of the fittest variant are less fit than $m$, so $p(L) = p(x_{\text{H}} < m)^{n-n_{\text{L}}}$ and likewise $p(H) = p(x_{\text{L}} < m)^{n_{\text{L}}}$, where $x_{\text{L}}$ is the fitness of an arbitrary mutant of the less-fit variant and $x_{\text{H}}$ is the fitness of an arbitrary mutant of the fittest variant. If $L$, any of the $n_{\text{L}}$ mutants can be the fittest as long as it is fitter than the remaining $n_{\text{L}} - 1$. Thus,

$$p(M = m|\lambda_{\text{L}}, L) = n_{\text{L}} p(x_{\text{L}} = m) p(x_{\text{L}} < m)^{n_{\text{L}}-1}$$

and likewise

$$p(M = m|\lambda_{\text{H}}, H) = (n - n_{\text{L}}) p(x_{\text{H}} = m) p(x_{\text{H}} < m)^{n-n_{\text{L}}-1}.$$

According to the two-parent model, $(x_{\text{L}} + \Delta x) \sim \text{Exponential}(\lambda_{\text{L}})$, $x_{\text{H}} \sim \text{Exponential}(\lambda_{\text{H}})$, and $\lambda_{\text{L}}, \lambda_{\text{H}} \overset{\text{i.i.d.}}{\sim} \text{Exponential}(\alpha)$. Thus,

$$\begin{aligned} p(M = m) = \alpha^2 \int_0^\infty \int_0^\infty & [n_{\text{L}} \lambda_{\text{L}} e^{-\lambda_{\text{L}}(m+\Delta x)} (1 - e^{-\lambda_{\text{L}}(m+\Delta x)})^{n_{\text{L}}-1} (1 - e^{-\lambda_{\text{H}} m})^{n-n_{\text{L}}} \\ & + (n - n_{\text{L}}) \lambda_{\text{H}} e^{-\lambda_{\text{H}} m} (1 - e^{-\lambda_{\text{H}} m})^{n-n_{\text{L}}-1} (1 - e^{-\lambda_{\text{L}}(m+\Delta x)})^{n_{\text{L}}}] \\ & \qquad e^{-\alpha(\lambda_{\text{L}}+\lambda_{\text{H}})} d\lambda_{\text{L}} d\lambda_{\text{H}}. \end{aligned}$$

Using Mathematica, we solve the integral $\text{E}(\log M) = \int_0^\infty p(M = m) \log(m) dm$ setting $n_{\text{L}} = 0$ and $n_{\text{L}} = n/2$ for various even $n$, and find the intersection.

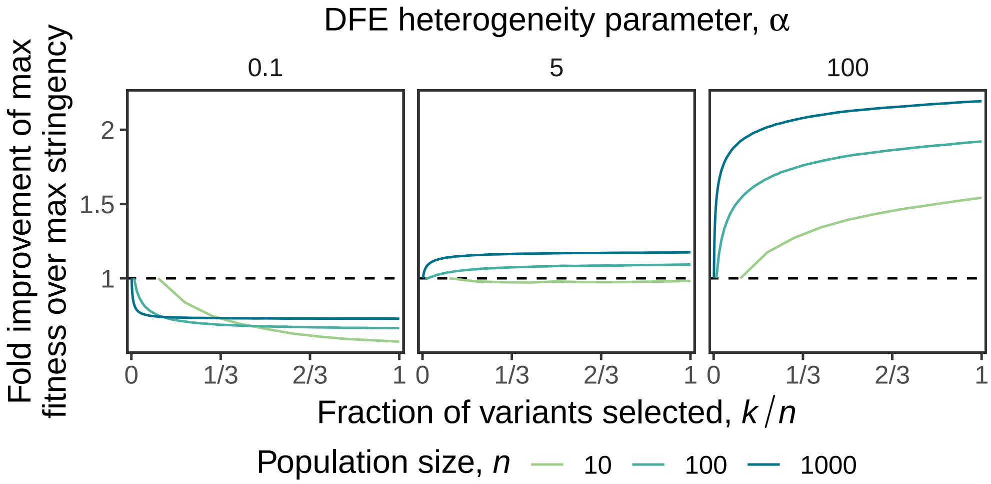


**Supplementary Figure 1. The immediate effect of selection stringency on fitness in an alternative $k$-parent model.** As in Fig 3B, but the DFE of the $i$th parent is Normal $\left(\sqrt{\lambda_i}, 1\right)$ where $\lambda_i \overset{\text{i.i.d.}}{\sim} \text{Exponential}(1/\alpha)$.

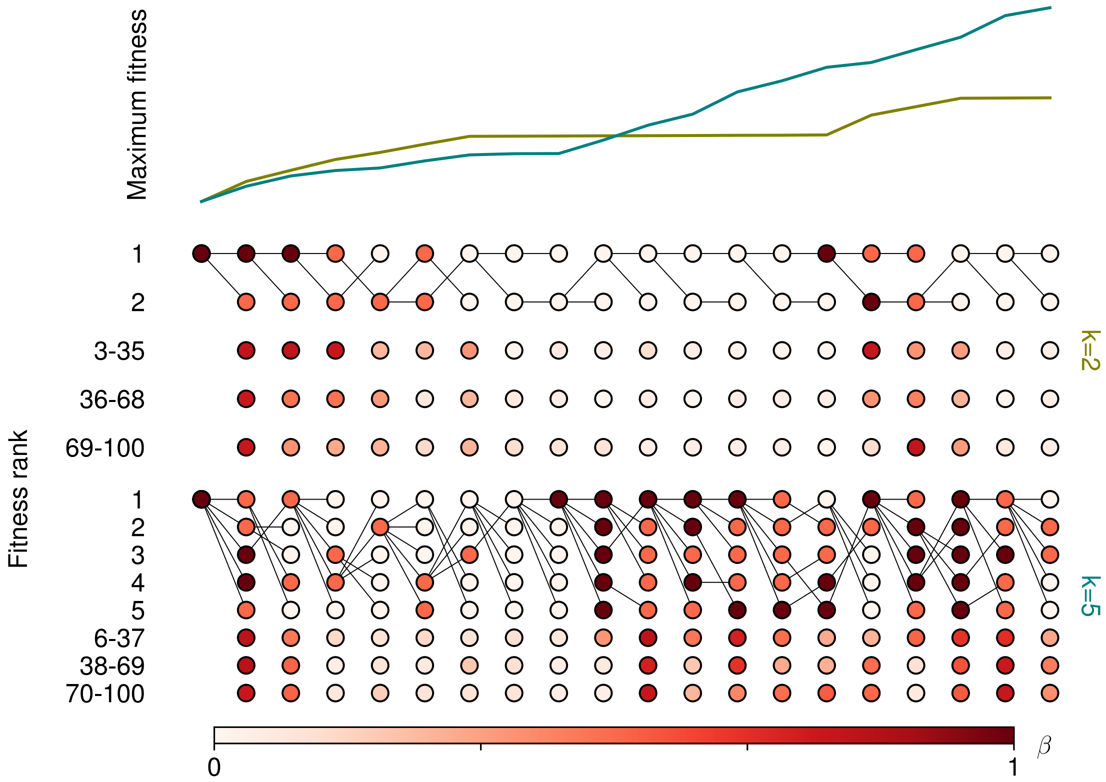


**Supplementary Figure 2. Longer-term dynamics of fitness with respect to selection stringency when DFE favorability can be recovered.** Example evolutions as in Fig 4A at two selection stringences, assuming $p = d = 1/2$ and a probability $1/20$ of recovering to $\beta = 1$.
